# Neural dynamics of associative learning during human sleep

**DOI:** 10.1101/372037

**Authors:** Andrés F. Canales-Johnson, Emiliano Merlo, Tristan A. Bekinschtein, Anat Arzi

## Abstract

Recent evidence indicate that humans can learn entirely new information during sleep. To elucidate the neural dynamics underlying sleep-learning we investigated brain activity during auditory-olfactory discriminatory associative learning in human sleep. We found that learning-related delta and sigma neural changes are involved in early acquisition stages, when new associations are being formed. In contrast, learning-related theta activity emerged in later stages of the learning process, after tone-odour associations were already established. These findings suggest that learning new associations during sleep is signalled by a dynamic interplay between slow-waves, sigma and theta activity.

## Introduction

The possibility to learn during sleep has intrigued humanity for over a century. In his 1911 science fiction novel “Ralph 124C 41+”, Hugo Gernsback described the *Hypnobioscope*, a device that transmits words directly to the sleeping brain such that they would be fully remembered in the next morning. However, decades of scientific efforts to teach sleeping humans new verbal information have been largely unsuccessful (for review (Peigneux et al. 2001)). Recently, the question of learning during sleep was revisited, and by applying simple forms of learning, such as associative and perceptual learning, it has been found that humans (Arzi et al. 2012, 2014; Ruch et al. 2014; Andrillon and Kouider 2016; Andrillon et al. 2017; Züst et al. 2019) and animals (De Lavilléon et al. 2015) can learn entirely new information during sleep. Yet, the brain mechanisms enabling learning of novel information during sleep are still unknown. Here, we aimed to identify brain processes supporting discriminatory associative learning during sleep.

We hypothesized that main brain sleep signals associated with consolidation and reactivation of information learned in awake state (Diekelmann and Born 2010; Oudiette and Paller 2013): slow-waves (delta: 0.5–4 Hz) (Marshall et al. 2006; Rasch et al. 2007; Antony et al. 2012), theta (4-7Hz) (Schreiner and Rasch 2015; Schreiner et al. 2015, 2018) and spindles (sigma: 11-16Hz) (Schabus et al. 2004; Tamminen et al. 2011; Laventure et al. 2016; Cairney et al. 2018), will be associated with learning novel associations during sleep. To test this hypothesis, we analysed electroencephalograph (EEG) activity recorded during auditory-olfactory partial-reinforcement conditioning in non-rapid eye movement (NREM) sleep (Figure 1a-d). On reinforced trials, each tone (400Hz or 1200Hz) was paired with either a pleasant or an unpleasant odour. Tone-odour pairings were counter-balanced across participants. On non-reinforced trials, either tone was presented without an ensuing odour, enabling the measure of learning-related neural correlates without the interference of odour. Stimuli were presented in blocks, each consisted of four reinforced trials and two non-reinforced trials (for detailed experimental design see methods, Figure S1 and Table S1). The conditioned response was the tone-induced sniff response, a behavioural change in nasal airflow in response to tone-odour pairings. Invariably, unpleasant odours drove smaller sniffs than pleasant odours (Arzi et al. 2012). Sleeping participants that learned these tone-odour associations subsequently showed modulated sniffs in response to tones alone, in accordance with the odour valence associated with the tone during sleep [behavioural data published in (Arzi et al. 2012). We found that learning-related delta and sigma activity are involved at early acquisition stages of the learning procedure, while learning-related theta activity emerges only after the discrimination is established, suggesting that timely modulation of slow-waves, sigma and theta rhythms during learning in sleep may prompt the encoding and stabilisation of new associative memories.

**Figure 1:**
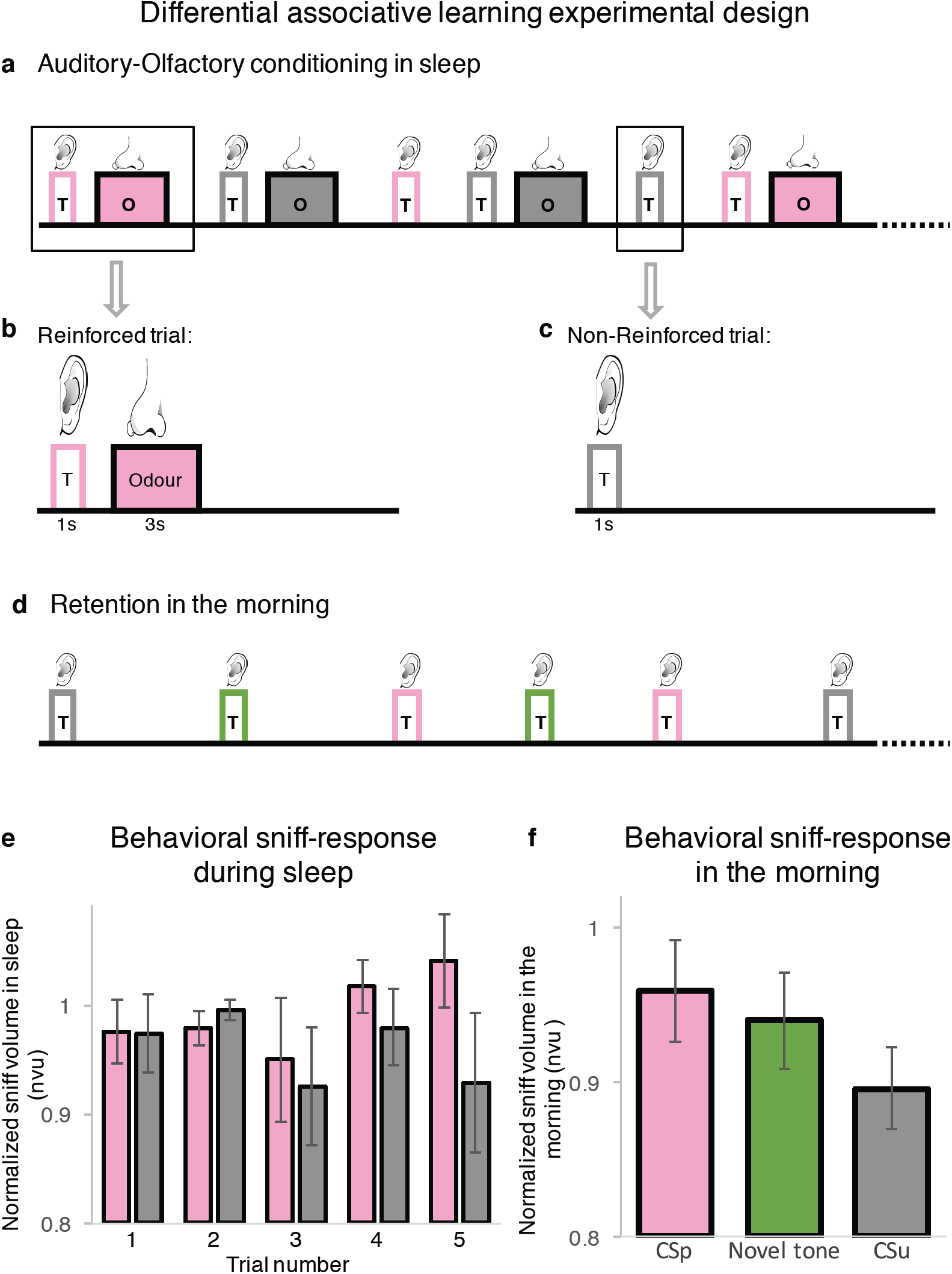
Auditory-olfactory discriminatory learning paradigm. **(a) Experimental design:** stimuli were generated in blocks of six trials: two reinforced trials with pleasant odour (pink), two reinforced trials with unpleasant odour (grey) and two non-reinforced trials (tone alone), one of each tone (see methods). T, tone; O, Odour. **(b)** On reinforced trials, each auditory stimulus (1,200 Hz or 400 Hz) was paired with either a pleasant (shampoo or deodorant) or unpleasant (rotten fish or carrion) odour. **(c)** On non-reinforced trials, either tone was presented alone. **(d)** Structure of the retention session performed during the subsequent morning where three auditory stimuli [1,200 Hz, 400 Hz and a novel 800-Hz tone (green), eight repetitions each] were presented without odours. **(e)** Normalized sniff response across continuous repetitions of a tone alone previously paired during sleep with a pleasant odour (CSp, pink) and continuous repetitions of a tone alone previously paired during sleep with unpleasant odour (CSu, grey) during the first five non-reinforced presentation of each CS during sleep, or first arousal, whichever came first. **(f)** Normalized sniff response during the retention session for CSp (pink bar), novel tone (800Hz, green bar), and for CSu (grey bar). nvu: <u>n</u>ormalized sniff <u>v</u>olume <u>u</u>nits [sniff volume divided by the baseline nasal inhalation (see Methods)]. The data used here was collected as part of a study that examined whether humans can learn new associations during sleep and was published independently (Arzi et al. 2012).

## Materials and Methods

The data used here was collected as part of a study that examined whether humans can learn new associations during sleep and was published independently (Arzi et al. 2012). Thus, detailed information about participants, experimental design, data acquisition and behavioural results can be found in the original article.

### Participants

Fourth-three healthy participants (mean age = 25.2 ± 3.2 years, 17 females) gave informed consent to procedures approved by the Weizmann Institute Ethics Committee. Participant exclusion criteria included use of medication, history of sleep disorders and nasal insults, or insufficient sleeping time. Out of these, 28 participants were presented with the auditory-olfactory conditioning during both NREM and REM sleep, and 15 participants during NREM sleep-only. Participants were unaware of specific experimental aims and conditions.

### Stimuli

Pleasant (shampoo or deodorant) and unpleasant (rotten fish or carrion) odorants were presented in a nasal mask by computer-controlled air-dilution olfactometer from an adjacent room (stimulus duration = 3 s, constant flow = 6 litres per minute). Tones (400, 800 and 1,200 Hz, duration = 1 s, at a non-arousing 40 dB) were presented by a loudspeaker ∼2 m from participants’ heads.

### Procedures

Participants arrived at the sleep laboratory at a self-selected time, based on their usual sleep pattern, typically at 11:00 pm. After fitting of the polysomnography devices (Iber et al. 2007), subjects were left alone in the darkened room to be observed from the neighbouring control room via infrared video camera. The experimenters observed the real-time polysomnography reading and, after they determined that the subject had entered the desirable sleep stage, they initiated the experimental protocol. The conditioned and unconditioned stimuli were partially reinforced at a ratio of 2:1; on reinforced trials (two-thirds of trials), each 1-s auditory conditioned stimulus (either 1,200 Hz or 400 Hz) was triggered by inhalation and paired with a 3-s olfactory unconditioned stimulus (either pleasant or unpleasant). On non-reinforced trials (one-third of trials), a tone triggered by inhalation was generated without an odorant (tone alone). Stimuli were generated in blocks of six trials (two reinforced trials with pleasant odour, two with unpleasant odour and two non-reinforced trials, one of each tone, randomized between blocks, Inter-trial interval 25-40 seconds). Tone-odour contingencies were counter-balanced across participants. The conditioned response was measured by the sniff response magnitude elicited during tone alone trials. During wakefulness the sniff response can be conditioned to a tone such that different tones can drive different sniffs (Resnik et al. 2011). Therefore, the sniff response was chosen to be the conditioned response in this experiment. In the NREM and REM group (28 participants), in a night without arousals/wakes within a window of 30 s from tone onset, five blocks were presented in NREM sleep, then the procedure was halted up to stable REM sleep, at which point an additional five blocks were presented. In NREM-only, the procedure was triggered during NREM sleep only (15 participants). If an arousal/wake was detected in the ongoing polysomnographic recording, the experiment was immediately stopped until stable sleep was resumed and continued up to a maximum of 18 blocks. Thus, the distribution between NREM and REM depended on each participant’s sleep structure. Since the experiment was halted following arousal or wake, different participants had different numbers of trials (Mean = 62.9 ± 19.3 trials) and varying training intervals between blocks imposed by individual sleep structure. About half an hour after spontaneous morning wake, conditioned response was tested in a retention procedure: three auditory stimuli, 1,200 Hz and 400 Hz that were presented during the night, and a new 800-Hz tone (eight repetitions each), were sequentially presented while nasal respiration was recorded. Retention procedure data from two subjects was lost due to technical problems.

### Polysomnography

Sleep was recorded by standard polysomnography (Iber et al. 2007). EEG (obtained from C3 and C4, referenced to opposite mastoid), electro-oculogram (placed 1 cm above or below and laterally of each eye, referenced to opposite mastoid), electromyogram (located bilaterally adjacent to the *submentalis* muscles), and respiration were simultaneously recorded (Power-Lab 16SP and Octal Bio Amp ML138, ADInstruments) at 1 kHz. Nasal respiration was measured using a spirometer (ML141, ADInstruments) and high-sensitivity pneumotachometer (#4719, Hans Rudolph) in line with the vent ports of the nasal mask.

### Nasal airflow analysis

Nasal inhalation volume in the retention session was normalised by dividing the sniff volume for each tone by the baseline nasal inhalation volume (averaged volume of 15 nasal inhalations preceding retention procedure onset). Participants’ normalized sniff volume differing by 3 SD were excluded (one participant).

### EEG analysis

EEG activity was recorded from C3 and C4 electrodes. Trials with EEG artefacts within a 5.5-second window before tone onset or 5-second after tone onset were excluded. EEG spectral analysis in the 0.5-40 Hz frequency range for all non-reinforced trials that met study criteria was conducted using Hilbert transform on a 10.5-second window using customized MATLAB scripts, with time-frequency resolution of 0.5Hz bin per 1 msec. The data was filtered using Hamming window sinc finite impulse response (FIR) filter implemented in EEGLAB. The filter order/transition band width is 25% of lower bandpass edge but not lower than 2Hz where possible. Specifically, Hilbert transform was conducted on a longer time window to avoid edge artefact and the time points before and after the 10.5-second window of interest were trimmed. The power in each trial was then z-scored. Non-reinforced trials from each participant were averaged across the two electrodes, and then averaged across trials to create a single time-frequency representation per participant, and per condition for the early-training phase (averaged across the non-reinforced trials in the first five blocks) and late-training phase (averaged across the non-reinforced trials in the sixth to the last blocks) during NREM sleep.

### Inclusion/exclusion criteria

An independent experienced sleep technician, blind to experimental conditions and to stimulus onset/offset times, scored the data off-line according to American Academy of Sleep Medicine criteria (Iber et al. 2007). We then used these blindly obtained scorings to include participants and/or trials. We included only EEG artefact-free trials without wake or arousal within 30 s of tone onset presented during NREM sleep. Importantly, trials presented in REM sleep in the ‘NREM and REM’ group where not included in the analysis. To avoid a bias of the results by individual trials we included in the analysis only participants with a minimum of 10 EEG-clean and arousals-free non-reinforced trials. Out of the 43 participants, five participants had less than 10 EEG-clean and arousals-free non-reinforced trials. Excluding these five participants, data from 38 participants remained for the EEG power analysis. In addition, two participants lacked the retention paradigm (morning testing) due to technical error and one participant’s sniff response was an outlier (> 3 SD). Excluding these three participants, data from 35 participants remained for the regression analyses presented in the supplementary materials. Total number of included non-reinforced trials in NREM sleep per learning phase and condition were 153 trials for tone alone previously paired with an unpleasant odour (CSu) and 170 trials for tone alone previously paired with a pleasant odour (CSp) in early-training, and 146 trials for CSu and 133 for CSp for late-training. The average number of included trials per participant was 4.03 ± 0.94 for CSu and 4.47 ± 0.98 for CSp in early-training, and 3.8 ± 2.9 for CSu and 3.5 ± 2.7 for CSp in late-training. The average number of trials was similar between early- and late-training phases however the variance was larger in the late-training due to individual differences in sleep architecture between participants.

### Statistical analysis

A permutation-based statistical test of time-frequency data was applied using FieldTrip (Oostenveld et al. 2011) and customized scripts. Time-frequency representation of each condition (CSu and CSp) and learning phase (early and late) in NREM sleep, were submitted to a cluster-based non-parametric permutation test (Maris and Oostenveld 2007), to determine in which time-frequency points in a 5-second window from tone onset there was a significant (α = 0.05) change from baseline (−5500 ms to −500 ms pre-tone onset; Figure 2a-d). In addition, the power envelope in each frequency band of interest [delta (0.5-4 Hz), theta (4-7 Hz) and sigma (11-16Hz)] was calculated for each condition separately (CSu and CSp), and submitted to cluster-based non-parametric permutation test to determine in which time points in a 5-second window from tone onset there was a significant increase from baseline per condition per learning phase. To determine whether there was a significant difference in power between CSu and CSp, non-reinforced trials were averaged across conditions (CSu and CSp) and learning phases (early and late). Then the averaged signal was submitted to cluster-based non-parametric permutation test to identify the time points where increase in power following the conditioned stimuli was significantly greater than baseline. A significant cluster was found in each one of the three frequency bands: delta (13 - 2184 ms, p_cluster_ < 0.001), theta (1 - 850 ms, p_cluster_ < 0.001) and sigma (727 - 1553 ms, p_cluster_ < 0.001). The cluster-based non-parametric permutation test between CSu and CSp in each learning phase was applied on the time window defined by the above-mentioned clusters in each frequency band. Multiple regression and correlation analyses were performed using Matlab and open-source statistical software JASP (JASP Team (2017), version 0.8.3.1). Nonparametric effect size was calculated by the following formula r = z/sqrt(n) (Rosenthal et al. 1994).

**Figure 2:**
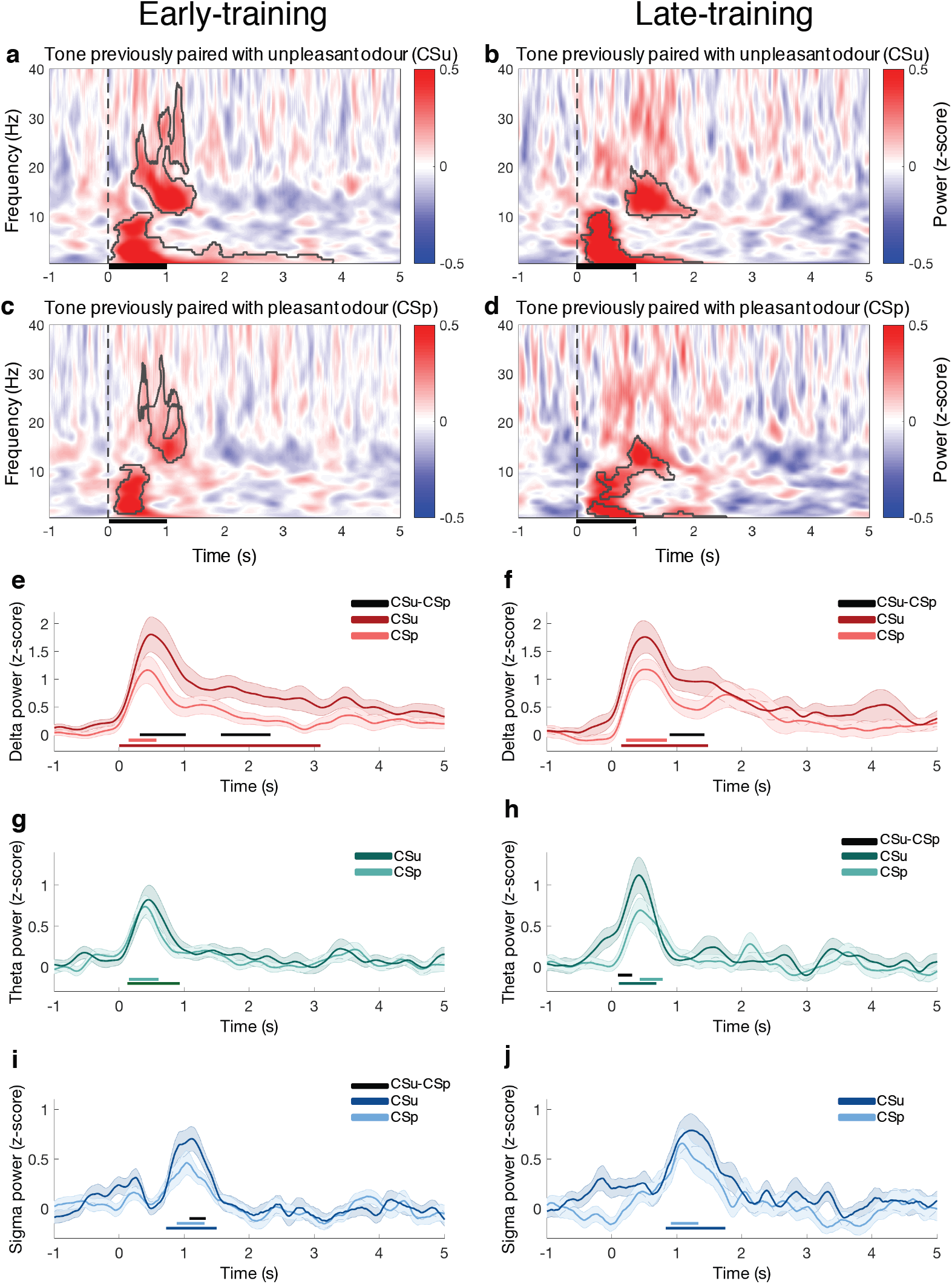
Learning-related electrophysiological activity in non-reinforced trials during NREM sleep. **(a-d)** Time–frequency decomposition of the EEG signal averaged across C3 and C4 electrodes in non-reinforced trials during NREM sleep time-locked to: **(a-b)** tone previously paired during sleep with an unpleasant odour (CSu) during **(a)** early-training phase, or **(b)** late-training phase; **(c-d)** tone previously paired during sleep with a pleasant odour (CSp) during **(c)** early-training phase, or **(d)** late-training phase. Areas inside black contours indicate significant deviations from zero compared to baseline (cluster permutation *t*-test, *P* _cluster_ < 0.05), each dotted vertical line represents tone onset, tone duration was 1 sec. EEG delta power during **(e)** early-training and **(f)** late-training for CSu (dark red) and CSp (light red). EEG theta power during **(g)** early-training and **(h)** late-training for CSu (dark green) and CSp (light green). EEG sigma power during **(i)** early-training and **(j)** late-training for CSu (dark blue) and CSp (light blue). Horizontal lines in the colour of the curve represent significant deviation from zero compared to baseline. Horizontal black lines represent significant difference between CSu and CSp (cluster permutation test, *P* _cluster_ < 0.05).

## Results

First, we verified that the behavioural results observed in the current examined NREM sleep dataset (see Methods) were similar to those reported before (The data was collected as part of a previous study published independently; Arzi et al. 2012). In order to investigate learning dynamics during discriminatory associative learning in NREM sleep we first examined the behavioural sniff response dynamics. We extracted the sniff volume during NREM sleep for a tone alone previously paired during sleep with an unpleasant odour (CSu) and for a tone alone previously paired during sleep with a pleasant odour (CSp) up to the fifth non-reinforced trial of each condition or first arousal, whichever came first (see Methods). One can see a gradual increase in CSp-CSu sniff volume difference across trials (Figure 1e). Moreover, the CSp-CSu sniff volume difference was significantly larger in the 4th - 5th block in comparison to the 1st - 2nd block (p < 0.05), similarly to what was previously shown in a subset of the dataset (Arzi et al. 2012). These findings suggest that the new tone-odour discriminatory associations were learned within the first five blocks. Therefore, we analysed the effects of training during sleep in early- (first five blocks; where discriminatory paired associations were acquired) and late-training (sixth to the last block (M = 5.5 ± 3.2 SD); where paired associations were well established). The following morning, during a test session, we observed that sniff response was larger for CSu than for CSp (*t*_35_ = 2.86, p = 0.015; excluding outlier: *t*_34_ = 2.75, p = 0.017; Figure S3c), suggesting the new associations learned during sleep were stored as memories readily retrievable upon awake (Figure 1f).

Then, to elucidate discriminatory associative learning-related brain dynamics during sleep, we analysed brain activity during non-reinforced CSu and CSp trials during NREM sleep for early- and late-training phases separately. Analysis of delta frequency band showed that during both early- and late-training, CSu elicited higher delta power in comparison to CSp (early-training: cluster1 396-1096 msec, p = 0.013, effect size r = 0.60; cluster2 1551-2320 msec, p = 0.011, effect size r = 0.43; Figure 2a,c,e; late-training: cluster 888-1418 msec, p = 0.028, effect size r = 0.42; Figure 2b,d,f), suggesting a learning-related delta modulation. Interestingly, the discriminatory neural response was different between the two training phases. While in the early-training phase two clusters were revealed, in the late-training phase only one cluster was found. While there was overlapping between the first cluster in early-training and the cluster in late-training, the second cluster from early-training uncovered a prolonged learning-related differential response that was not observed in the late-training phase. Furthermore, the differential response between CSu and CSp in early and late-training was similar in the first 1500 msec bout (no cluster) but was larger in early-training compared to late-training in the second 1500 msec bout from tone onset (1959-2273 msec, p_cluster_ = 0.05, effect size r = 0.31). These findings indicate a modulation of delta activity between early- and late-training, suggesting this neural correlate could signal the acquisition of new associative memory traces.

Similar analysis of theta frequency band revealed an opposite response pattern to the one observed in delta. No cluster was found in theta power when comparing CSu and CSp in early-training (Figure 2a, c, g), while late-training CSu elicited higher theta power (cluster 94-311 msec, p = 0.026, effect size r = 0.36; Figure 2b, d, h) compared to CSp. Moreover, the differential neural response between CSu and CSp was larger in late-training when compared with early-training (cluster = 1-156, msec, p = 0.05, effect size r = 0.37). These findings suggest that learning-related theta modulations emerge when new associations are well-trained or already established.

Analysis of sigma frequency band showed that CSu trials induced higher power than CSp homologous events in early-training (cluster 1080-1321 msec, p = 0.032, effect size r = 0.32; Figure 2a, c, i), but not in late-training (no cluster) (Figure 2b, d, j). However, we did not find reliable differences between early- and late-training in CSu and CSp differential response (no cluster). Thus, modulation in sigma power may underlie acquisition of associative memories in early learning stages, but may not have a distinct contribution to early-versus late-training stages.

## Discussion

Here, we aimed to elucidate the brain activity supporting discriminatory associative learning in sleep. Using EEG recordings during auditory-olfactory conditioning we uncovered learning-related delta, sigma and theta power modulation in NREM sleep. Moreover, in the course of discriminatory associative learning, learning-related delta and sigma activity are modulated at early acquisition stages while theta activity modulation emerges only after stimuli discrimination is well established. These effects were evident despite the variability in training history introduced by individual differences in sleep architecture (Table S1 and Figure S1).

During the discriminatory associative learning procedure sleeping participants learned to implicitly discriminate between tones predicting odours with different valence. This process involves learning two independent contingencies between specific tones and odours, and results in the ability to discriminate between the expected value of each tone. During the early-training phase, spanning the first five training blocks, the new tone-odour associations were readily acquired as indicated by the behavioural sniff response (Figure 1e). However, as contingency learning and discrimination occur during the same time frame, and discrimination is an integral part of the learning, the observed learning-related EEG power may reflect either or both of these processes. If a specific brain activity correlate is involved in contingency encoding it should be more apparent when prediction error is high (i.e. during early-training phase). On the contrary, if the same brain activity correlate indicates stimuli discrimination it should be increasingly recruited or stay constant as training progress from early to late trials. Thus, learning-related delta power modulation observed in cluster1 in both early- and late-training suggests an involvement of slow waves in the discrimination process occurring all along training. Modulation of delta power in cluster2 and sigma power observed specifically in early-training implies a role of these brain correlates in acquisition of new associations during sleep. Notably, the contribution of sleep depth to learning cannot be fully dissociated from training phase. In late-training, trials distribution was even between N2 and N3, however in early-training the vast majority of trials were presented in N3 (supplementary materials and Table S2), implying that sleep depth may interact with different stages of the learning process. Altogether, these findings suggest that slow-waves and spindles activity is part of the required conditions for encoding of novel associations in sleep.

To date, only a few studies investigate the neural activity underlying sleep-learning (Peigneux et al. 2001; Ruch et al. 2014; Andrillon and Kouider 2016; Andrillon et al. 2017; Farthouat et al. 2018; Züst et al. 2019). The observed learning-related delta power modulation in this study is in line with recent findings showing that successful verbal associative learning during NREM sleep is bound to slow-wave peaks (Ruch et al. 2014; Züst et al. 2019), and that unsuccessful retrieval of auditory perceptual learning during NREM sleep is associated with decreased delta power (Andrillon et al. 2017). In addition, the absence of theta modulation by learning during early-training stage is in agreement with findings of low theta activity during vocabulary encoding in NREM sleep (Züst et al. 2019), and could imply that theta activity does not signal encoding of new associations in NREM sleep. That said, perceptual learning during REM sleep was found to increase theta power (Andrillon et al. 2017). On the other hand, learning-related theta activity in later phases of the training procedure, after the new associations were formed, may be involved in strengthening already-established memories. For sigma, a more complex picture emerges. We observed learning-related modulation in early-training, however not in late-training and found no interaction between learning phases. Other studies, found no relationship between sigma activity and associative learning retrieval (Züst et al. 2019), and marginal modulation during perceptual learning (Andrillon et al. 2017). Taken together, these findings suggest that the same brain rhythm may have different roles during different learning processes and sleep stages.

Delta, sigma and theta activity are key players in memory reactivation and consolidation of information learned before sleep (Diekelmann and Born 2010; Oudiette and Paller 2013). Slow-waves have been associated with and causally related to memory consolidation (Marshall et al. 2006; Rasch et al. 2007; Antony et al. 2012); spindles have a role in integrating new memories and existing knowledge, and in memory consolidation (Diekelmann and Born 2010; Tamminen et al. 2011; Antony et al. 2012, 2019; Oudiette and Paller 2013; Cairney et al. 2018); theta is involved in memory consolidation and reactivation of information learned while awake (Schreiner and Rasch 2015; Schreiner et al. 2015, 2018). Unlike during sleep-learning, the involvement of these brain oscillations in memory formation, consolidation and retrieval is well-established (Diekelmann and Born 2010; Hanslmayr et al. 2016; Antony et al. 2019). However, whether consolidation, targeted memory reactivation and sleep-learning share similar brain mechanism is still an open question.

Understanding the brain process underlying acquisition, consolidation and retrieval of new information presented during sleep constitutes an important step in the course of identifying what and how is possible to learn during sleep (Simon and Emmons 1956; Wood et al. 1992; Arzi et al. 2012, 2014; Ruch et al. 2014; Andrillon and Kouider 2016; Andrillon et al. 2017; Farthouat et al. 2018; Züst et al. 2019). Here, we start to elucidate part of these processes showing that encoding of a discriminatory associative memory during sleep is associated with a dynamic interplay between learning-related slow-waves, sigma and theta activity. Timely modulation of these brain rhythms occurring during learning in sleep may determine the acquisition and storage of new associative memories.

## Acknowledgements

This work was supported by the Blavatnik family Foundation [to AA], Royal Society – Kohn International fellowship [NF150851 to AA], European Molecular Biology Organization (EMBO) fellowship [ALTF 33-2016 to AA], Fondo Nacional de Desarrollo Científico y Tecnológico [FONDECYT N 1171200 to ACJ], and The Wellcome Trust [WT093811MA to TAB]. Collection of the data at the Weizmann Institute of Science in Israel was supported by the Rob and Cheryl McEwen Fund for Brain Research. We are indebted to Noam Sobel for his valuable contribution to this work. We are grateful to Sridhar R. Jagannathan, Daniel Bor, Roni Kahana-Zweig, and Lavi Secundo for their comments and for fruitful discussions.

## Author Contributions

Conceptualization: AFC, EM, TAB, AA; Methodology: AFC, EM, TAB, AA Software: AFC, AA; Validation: AA; Formal analysis: AFC, AA; Investigation: AA Resources: TAB, AA; Data curation: AA; Writing - original draft: AA; Writing – Review & editing: AFC, EM, TAB, AA; Visualization: AFC, EM, TAB, AA; Supervision: AA, TAB; Project Administration: AA; Funding Acquisition: AFC, TAB, AA.

## Declaration of Interests

The authors declare no competing interests.

## Data availability

The raw data of this manuscript are available for download at: https://www.weizmann.ac.il/neurobiology/worg/materials.html.

