## Supplementary materials for "Neural dynamics of associative learning during human sleep"

for

[1]Department of Psychology, University of Cambridge, Cambridge, CB2 3EB UK. [2]Center for Social and Cognitive Neuroscience (CSCN), Universidad Adolfo Ibanez, Santiago, Chile. [3]The Neuropsychology and Cognitive Neurosciences Research Center (CINPSI Neurocog), Universidad Católica del Maule, Talca, Chile. [4]IFIBIO-Houssay, Facultad de Medicina, Universidad de Buenos Aires - CONICET, Buenos Aires, Argentina. [5]School of Psychology, University of Sussex, Brighton BN1 9RH, UK

*To whom correspondence should be addressed:

Anat Arzi

Department of Psychology

University of Cambridge

Cambridge, CB2 3EB, UK

Tel: (+) 44 1223765289

**Associative learning experimental procedure**

Sleep architecture dictated the structure of auditory-olfactory conditioning training procedure. The training procedure was initiated once the participant entered stable sleep and was based on the individual sleep structure. The individual differences in sleep architecture introduced variability in the training procedure structure between participants. Figure S1 and Table S1 detail the duration of the early-training, late-training, total training procedure, and the temporal gap between early- and late-training.

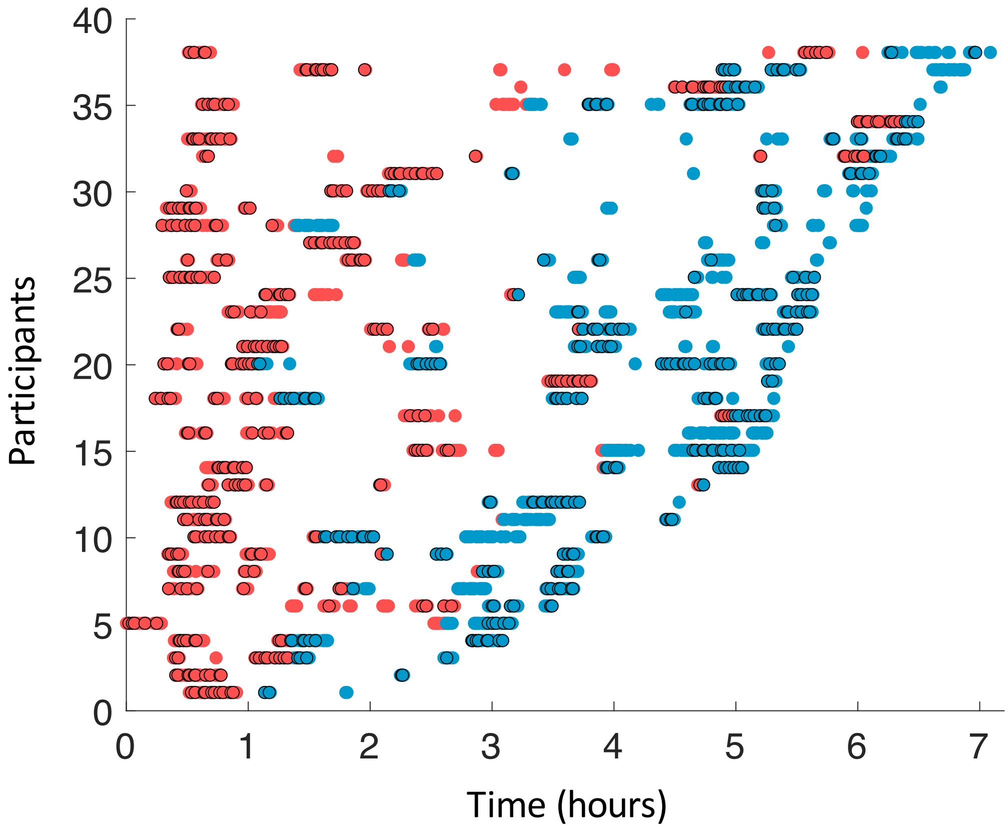


**Figure S1: Auditory-olfactory discriminatory conditioning training procedure** Temporal structure of the training procedure was based on individual sleep architecture. Each dot represents a trial, during early-training (red) or late-training (blue), and each row is a participant. Dots with black edge represent the non-reinforced trials in NREM sleep included in the analysis.

**Table S1: Auditory-olfactory discriminatory conditioning training records**

| Subject number | Early-training duration | Late-training duration | Gap between early and late-training | Total training duration |
|---|---|---|---|---|
| 1 | 00:23:18.1 | 00:40:30.1 | 00:13:49.1 | 01:17:37.1 |
| 2 | 00:22:17.1 | 00:01:39.1 | 01:28:13.1 | 01:52:09.1 |
| 3 | 00:57:18.1 | 01:18:01.1 | 00:02:11.1 | 02:17:30.1 |
| 4 | 00:57:15.1 | 01:43:46.1 | 00:00:33.1 | 02:41:33.1 |
| 5 | 02:36:27.1 | 00:32:52.1 | 00:00:30.1 | 03:09:49.1 |
| 6 | 01:20:16.1 | 00:31:17.1 | 00:16:42.1 | 02:08:15.1 |
| 7 | 01:17:40.1 | 01:49:13.1 | 00:00:29.1 | 03:07:22.1 |
| 8 | 02:30:02.1 | 00:46:54.1 | 00:02:03.1 | 03:18:58.1 |
| 9 | 01:45:01.1 | 01:36:31.1 | 00:00:31.1 | 03:22:03.1 |
| 10 | 01:01:23.1 | 02:20:43.1 | 00:00:34.1 | 03:22:39.1 |
| 11 | 02:36:46.1 | 01:23:52.1 | 00:00:51.1 | 04:01:29.1 |
| 12 | 00:21:10.1 | 01:34:00.1 | 02:14:55.1 | 04:10:05.1 |
| 13 | 04:02:51.1 | 00:00:31.1 | 00:00:43.1 | 04:04:05.1 |
| 14 | 03:16:01.1 | 01:07:44.1 | 00:00:33.1 | 04:24:18.1 |
| 15 | 01:33:24.1 | 01:12:21.1 | 00:01:49.1 | 02:47:35.1 |
| 16 | 00:47:49.1 | 00:38:42.1 | 02:36:44.1 | 04:03:15.1 |
| 17 | 02:41:34.1 | 00:18:43.1 | 00:01:20.1 | 03:01:37.1 |
| 18 | 01:00:53.1 | 04:02:10.1 | 00:00:49.1 | 05:03:53.1 |
| 19 | 00:21:30.1 | 00:04:08.1 | 01:26:40.1 | 01:52:17.1 |
| 20 | 00:46:09.1 | 04:16:10.1 | 00:00:30.1 | 05:02:49.1 |
| 21 | 01:12:05.1 | 03:07:04.1 | 00:09:21.1 | 04:28:30.1 |
| 22 | 03:18:04.1 | 01:46:32.1 | 00:00:33.1 | 05:05:09.1 |
| 23 | 00:26:10.1 | 02:06:10.1 | 02:15:07.1 | 04:47:28.1 |
| 24 | 02:02:49.1 | 02:26:17.1 | 00:00:34.1 | 04:29:41.1 |
| 25 | 00:22:07.1 | 01:58:25.1 | 02:56:57.1 | 05:17:29.1 |
| 26 | 01:47:39.1 | 03:18:50.1 | 00:04:04.1 | 05:10:33.1 |
| 27 | 00:22:13.1 | 01:01:57.1 | 02:52:03.1 | 04:16:14.1 |
| 28 | 00:22:45.1 | 04:38:28.1 | 00:00:30.1 | 05:01:44.1 |
| 29 | 00:41:07.1 | 02:07:40.1 | 01:10:57.1 | 03:59:44.1 |
| 30 | 01:39:12.1 | 03:57:18.1 | 00:00:34.1 | 05:37:03.1 |
| 31 | 00:23:48.1 | 02:58:14.1 | 00:36:12.1 | 03:58:13.1 |
| 32 | 05:27:17.1 | 00:09:56.1 | 00:01:26.1 | 05:38:39.1 |
| 33 | 00:21:25.1 | 02:44:42.1 | 02:46:42.1 | 05:52:49.1 |
| 34 | 00:21:28.1 | 00:06:47.1 | 00:01:32.1 | 00:29:47.1 |
| 35 | 01:46:58.1 | 03:12:22.1 | 00:01:31.1 | 05:00:51.1 |
| 36 | 01:41:12.1 | 01:45:06.1 | 00:00:33.1 | 03:26:51.1 |
| 37 | 02:34:13.1 | 01:59:21.1 | 00:53:35.1 | 05:27:09.1 |
| 38 | 04:51:45.1 | 00:50:24.1 | 00:12:30.1 | 05:54:40.1 |

**Table S2: Trials distribution between N2 and N3 sleep**

| Subject number | Early-training | | | | Late-training | | | |
|---|---|---|---|---|---|---|---|---|
| | CSp | | CSu | | CSp | | CSu | |
| | N2 | N3 | N2 | N3 | N2 | N3 | N2 | N3 |
| 1 | 1 | 2 | 0 | 4 | 2 | 2 | 2 | 2 |
| 2 | 1 | 3 | 0 | 4 | 5 | 0 | 5 | 0 |
| 3 | 0 | 5 | 0 | 5 | 0 | 0 | 0 | 0 |
| 4 | 0 | 5 | 0 | 5 | 1 | 0 | 2 | 0 |
| 5 | 3 | 2 | 2 | 2 | 2 | 0 | 2 | 0 |
| 6 | 0 | 3 | 0 | 3 | 3 | 1 | 3 | 3 |
| 7 | 0 | 4 | 0 | 4 | 2 | 0 | 1 | 1 |
| 8 | 0 | 2 | 0 | 3 | 2 | 0 | 3 | 0 |
| 9 | 0 | 4 | 0 | 5 | 1 | 0 | 0 | 0 |
| 10 | 1 | 5 | 1 | 2 | 0 | 3 | 0 | 4 |
| 11 | 0 | 3 | 0 | 3 | 3 | 5 | 4 | 5 |
| 12 | 0 | 4 | 0 | 5 | 4 | 1 | 5 | 0 |
| 13 | 0 | 5 | 0 | 5 | 6 | 6 | 4 | 8 |
| 14 | 1 | 4 | 1 | 4 | 3 | 4 | 3 | 4 |
| 15 | 1 | 3 | 0 | 3 | 5 | 1 | 7 | 0 |
| 16 | 0 | 5 | 0 | 4 | 2 | 2 | 1 | 2 |
| 17 | 1 | 3 | 1 | 3 | 5 | 4 | 4 | 5 |
| 18 | 0 | 3 | 0 | 1 | 4 | 0 | 4 | 1 |
| 19 | 0 | 5 | 0 | 4 | 0 | 0 | 0 | 0 |
| 20 | 5 | 0 | 3 | 0 | 7 | 0 | 9 | 0 |
| 21 | 3 | 0 | 2 | 0 | 3 | 0 | 3 | 0 |
| 22 | 0 | 5 | 0 | 5 | 1 | 6 | 1 | 6 |
| 23 | 0 | 5 | 0 | 4 | 0 | 1 | 0 | 4 |
| 24 | 0 | 4 | 0 | 4 | 0 | 1 | 0 | 2 |
| 25 | 0 | 5 | 0 | 4 | 0 | 1 | 2 | 1 |
| 26 | 2 | 5 | 0 | 4 | 0 | 3 | 0 | 3 |
| 27 | 5 | 0 | 4 | 0 | 0 | 1 | 0 | 1 |
| 28 | 3 | 2 | 3 | 2 | 2 | 0 | 2 | 0 |
| 29 | 5 | 0 | 5 | 0 | 2 | 0 | 2 | 0 |
| 30 | 0 | 5 | 0 | 3 | 1 | 5 | 1 | 6 |
| 31 | 0 | 5 | 0 | 4 | 0 | 4 | 0 | 4 |
| 32 | 1 | 2 | 0 | 4 | 0 | 4 | 0 | 4 |
| 33 | 0 | 5 | 0 | 5 | 0 | 1 | 0 | 1 |
| 34 | 0 | 4 | 0 | 4 | 0 | 2 | 0 | 2 |
| 35 | 0 | 5 | 1 | 4 | 0 | 4 | 0 | 3 |
| 36 | 0 | 5 | 0 | 5 | 0 | 1 | 0 | 0 |
| 37 | 0 | 5 | 0 | 4 | 0 | 3 | 0 | 3 |
| 38 | 0 | 5 | 0 | 5 | 0 | 1 | 0 | 1 |
| **Total** | **33 (19%)** | **137 (81%)** | **23 (15%)** | **130 (85%)** | **66 (50%)** | **67 (50%)** | **70 (48%)** | **76 (52%)** |

**Table S3: Sleep parameters**

| Total sleep duration (min) | % WASO | % N1 | % N2 | % N3 | % REM |
|---|---|---|---|---|---|
| 319.59 ± 73.02 | 20.41 ± 10.04 | 3.03 ± 2.42 | 41.46 ± 12.86 | 18.04 ± 5.61 | 17.06 ± 7.04 |

N1, N2, N3: NREM sleep stages N1, N2, and N3, REM: rapid eye movement sleep, WASO: wake after sleep onset. Values are means ± SEM

**Learning-related behavioural responses**

To test whether the tone-induced sniff response reflects learning and not merely an orientation response, we conducted a control experiment published previously (Arzi et al. 2012): "To verify that previously unpaired tones alone do not elicit variable responses on the basis of tone frequency, we presented a control group with the retention protocol awake, without conditioning during sleep. We found no difference in nasal inhalation volume between 400- and 1,200-Hz tones (normalized volume: 400 Hz, 1.08 ± 0.17 nvu; 1,200 Hz, 1.11 ± 0.19 nvu; $t_9 = 0.72$, $P > 0.49$, $n = 10$; Fig. 3e)."

In the current work we also tested whether the control group collapsed CS+ (CSu + CSp) inhalation volume showed a reduction from baseline and found no change from baseline inhalation ($t_9 = 1.4$, $p = 0.2$), while the same analysis in the conditioning group showed a significant reduction from baseline ($t_{35} = 2.6$, $p = 0.015$). In addition, the sniff response to the novel 800Hz tone was intermediate compared with those from CSp and CSu (Normalized sniff volume CSu: 0.90 ± 0.16 nvu (normalized volume units), CSp: 0.96 ± 0.20 nvu, novel 800Hz tone: 0.94 ± 0.18 nvu; Figure 1f). CSu sniff response was greater (larger reduction or smaller normalized volume) than for the novel tone ($t_{35} = 1.97$, $p = 0.05$), while there was no difference between CSp sniff response and for novel tone ($t_{35} = 0.68$, $p = 0.5$). Altogether, these analyses demonstrate that modulation in nasal inhalation volumes following CS+ was learning-related and not a simple orientation response.

### Learning-related tone-evoked response during sleep

To test whether the learning-related differential response between CSu and CSp in the first 1500 msec merely reflects an evoked K-complex activity we compared the tone-evoked response between CSu and CSp and found no significant cluster in neither early- ($p_{cluster}$ = 0.13) nor late-training (no cluster) phases (Figure S2). These findings suggest that learning-related differential increase in delta power does not solely mirror the tone-evoked response.  In addition, to test whether there were differences in tone-evoked response between early and late sleep we compared the tone-evoked response between training phases by performing a cluster permutation test between early- and late-training on CSu and CSp separately, and found no significant cluster in tone-evoked response between early and late-training in neither CSp nor CSu (all $p's_{cluster}$ > 0.3), implying that tone-induced activity did not differ between training phases.

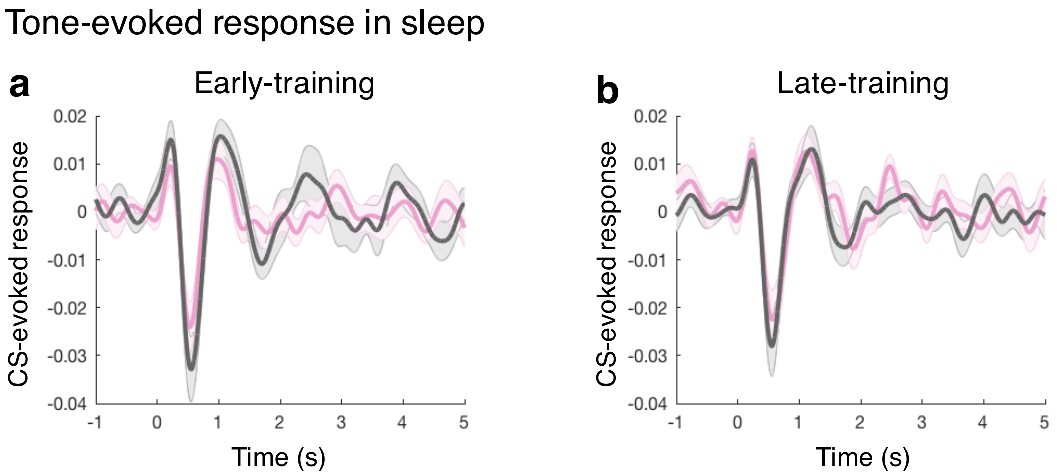


**Figure S2: Tone-evoked response during sleep**
**(a)** Tone-evoked response for CSp (pink) and CSu (grey) during early-training sleep.
**(b)** Tone-evoked response for CSp (pink) and CSu (grey) during late-training sleep.

**Relation between learning-related neural and behavioural responses**

We tested whether the observed learning-related brain correlates during sleep predict discriminatory memory evaluated during the following morning. As during training phase, the behavioural measurement of memory was the tone-induced sniff response, indexing odour expectation, and the learning-related brain correlate was tone-induced power. A multiple linear regression was calculated to predict CSu-CSp difference in sniff response based on CSu-CSp difference in power in the frequencies of interest: delta, sigma and theta. No significant relationship was found in either early- ($R^2$ = 0.01, p = 0.93) or late-training ($R^2$ = 0.08, p = 0.54). These findings suggest that the learning-related differential brain and learning-related differential behavioural responses are not directly link. However, the sniff response for CSu and CSp were highly correlated ($r$ = 0.57, p = 0.0004). Therefore, we averaged the sniff response and EEG power across CSu and CSp and calculated a multiple linear regression to predict the non-differential sniff response (tone-induced change in sniff volume) based on delta, sigma and theta power. We found a positive relationship between the sniff response in subsequent morning and EEG power in sleep during early- ($R^2$ = 0.32, p = 0.007) and late-training ($R^2$ = 0.25, p = 0.038). Early-training delta (B = 0.07, p = 0.037; Figure 3a) and sigma (B = 0.16, p = 0.009; Figure 3c), but not theta (B = -0.06, p = 0.38; Figure 3e), power predicted successful memory retrieval. Late-training theta (B = 0.12, p = 0.026; Figure 3f), but not delta (B = 0.015, p = 0.58; Figure 3b) nor sigma (B = -0.03, p = 0.51; Figure 3d), predicted successful memory retrieval. In other words, the greater the tone-induced EEG power during NREM sleep the larger the memory retention in the morning. Frequency bands modulated during learning in each training phase in sleep were the ones associated with memory retention in the subsequent morning. A possible interpretation for these findings is that the level of responsiveness to external stimuli presented during sleep is linked to the ability to learn new information associated with these stimuli and store it as memories.

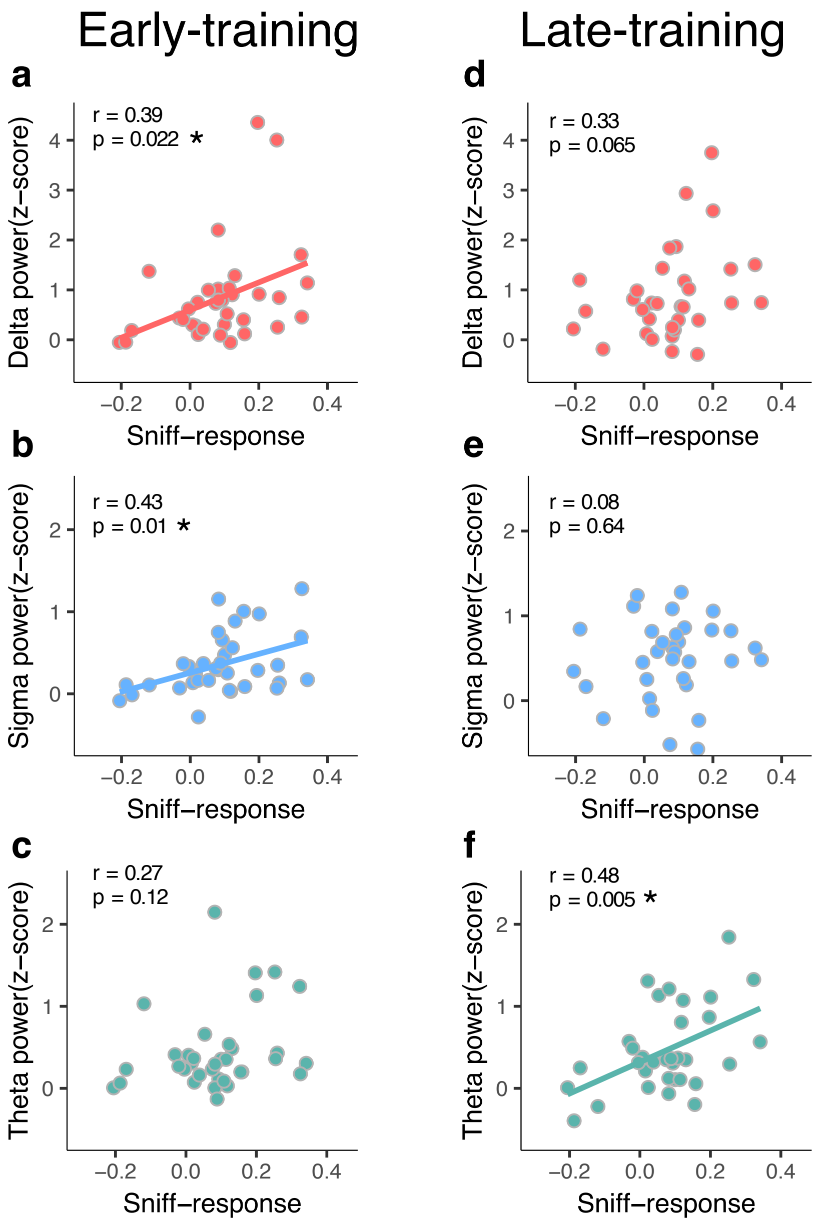


**Figure S3: Relation between learning-related EEG power in sleep and learning-related behavioural sniff response during test.** Correlation between tone-induced **(a)** delta (r = 0.39, p = 0.022), **(b)** sigma (r = 0.43, p = 0.01) and **(c)** theta (r = 0.27, p = 0.12) power during early-training in sleep and tone-induced sniff response in subsequent morning across CSu and CSp. Correlation between tone-induced **(d)** delta (r = 0.33, p = 0.065), **(e)** sigma (r = 0.08, p = 0.64) and **(f)** theta (r = 0.48, p =

0.005) power during late-training in sleep and tone-induced sniff response in subsequent morning across CSu and CSp. r denotes Pearson correlation coefficient, also see Supp. Figure S3; * denotes FDR-corrected significance threshold. The sniff response was calculated as 1 - normalized sniff volume.

*Training phases*

To directly compare the relationship between learning-related behavioural and brain responses in early- and late-training we conducted a linear regression analysis including both training phases in each frequency band separately. A multiple linear regression to predict the non-differential sniff response (tone-induced decrease in sniff volume) based on early- and late-training delta power revealed a positive relationship between the sniff response in subsequent morning and learning-related delta power in sleep ($R^2$ = 0.23, p = 0.02), where early-training delta (B = 0.05, p = 0.043;) but probably not late-training delta (B = 0.022, p = 0.39) power predicts memory retrieval. Explained variance was small and similar between early- and late-training, and no reliable difference was found between the two correlations (Fisher test for significance of the difference between two correlation coefficients: z = 0.27, p = 0.78). Similar analysis performed on theta frequency band revealed a positive relationship between sniff response in subsequent morning and theta power in sleep ($R^2$ = 0.15, p = 0.007), where early-training theta did not seem to predict memory retrieval (B = 0.048, p = 0.26;) and late-training theta did (B = 0.11, p = 0.01), however no significant difference was found between the two correlations (Fisher test: z = 0.97, p = 0.33). Last, the same multiple linear regression in sigma frequency band showed a possibly week association between sniff response in subsequent morning and sigma power in sleep ($R^2$ = 0.17, p = 0.06), with early-training sigma (B = 0.15, p = 0.025;) and most likely no late-training sigma (B = 0.023, p = 0.62) power predicting memory retrieval. No reliable difference was found between early- and late-training correlations (Fisher test: $z = 1.49$, $p = 0.14$).

*Sleep stage influence on learning*

In the main text, we analysed non-reinforced trials presented during NREM sleep. Yet in some of the participants the conditioning was presented also during REM sleep. Although not included in the analysis, conditioning in REM sleep in addition to conditioning in NREM sleep might have influenced learning, measured by the behavioural sniff response at test. We examined the sniff response in the 'NREM and REM conditioning' group, and in the 'NREM-only conditioning' group, and both groups have similar sniff response values and similar range (NREM and REM conditioning' group sniff response 0.063 ± 0.18; 'NREM-only conditioning' group 0.081 ± 0.12, Wilcoxon rank-sum test $p = 0.65$). The 'NREM and REM conditioning' group and 'NREM-only conditioning' group were colour coded for visualization (Figure S4).

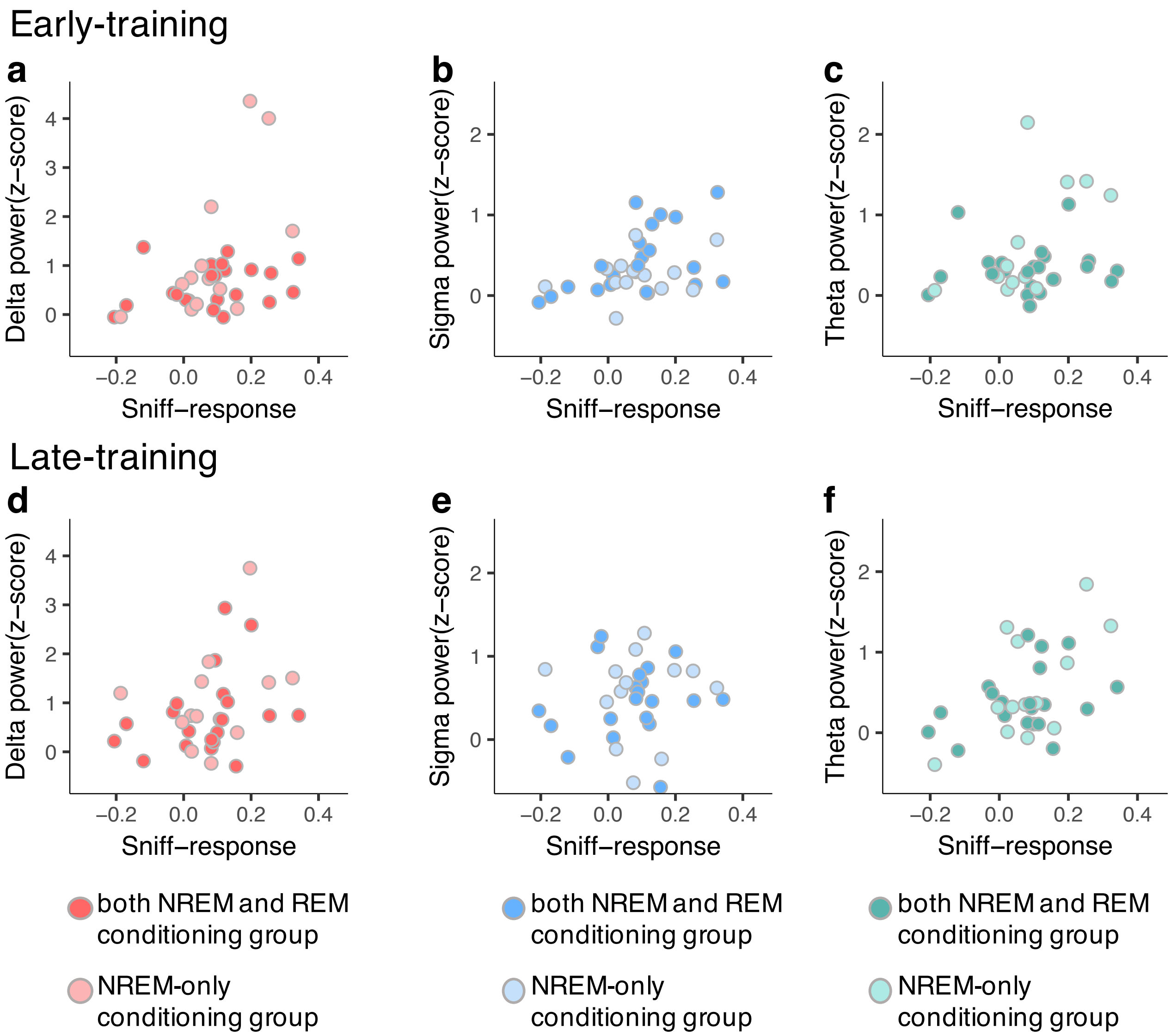


**Figure S4: Similar tone-induced EEG power and sniff response between conditioning in both NREM and REM sleep and NREM-only sleep**
**(a-c)** Relation between tone-induced **(a)** delta **(b)** sigma and **(c)** theta power during early-training in sleep and tone-induced sniff response in subsequent morning across CSu and CSp. **(d-f)** Relation between tone-induced **(d)** delta **(e)** sigma and **(f)** theta power during late-training in sleep and tone-induced sniff response in subsequent morning across CSu and CSp. Color-code: ‘both NREM and REM conditioning’ group: dark red (delta) dark blue (sigma) and dark green (theta); NREM-only conditioning’ group: light red (delta), light blue (sigma) and light green (theta).

*Correlations outliers*

One of the 36 participants showed a sniff response value (average of CSu and CSp combined) exceeding 3 SD from the mean of the group. Thus, in the main text we excluded that value when applying the linear regression and correlation analyses between learning-related EEG power in NREM sleep and sniff response in subsequent morning. When including the outlier value, the pattern of results remained constant: in early-training delta ($r$ = 0.34, p = 0.042) and sigma ($r$ = 0.37, p = 0.025), but not theta ($r$ = 0.23, p = 0.19) power positively correlated with the sniff response. During late-training, theta ($r$ = 0.36, p = 0.035), but not delta ($r$ = 0.25, p = 0.15) nor sigma ($r$ = 0.17, p = 0.32), power correlated with the sniff response.

*Sample size*

Sample size in sleep and memory studies usually ranges between 15 to 40, with most studies having ~20 participants per group. In our study the sample size was 38 and is considered in the higher end of the scale. However, due to individual sleep structure and EEG signal noise, in the late-training phase 4 participants had no non-reinforced trials, remaining 34 participants. We repeated the analysis for the early-training phase in the 34 participants and found the same results as with 38 participants (*34 participants*: delta early cluster 1: CSu = 1.54 normalised power, CSp = 0.76 normalised power, Wilcoxon sign rank test p = 0.00015, Delta early cluster 2: CSu = 0.83 normalised power, CSp = 0.27 normalised power, Wilcoxon sign rank test p = 0.0016, Sigma early cluster 1: CSu = 0.33 normalised power, CSp = 0.18 normalised power, Wilcoxon sign rank test p = 0.037; *38 participants*: delta early cluster 1: CSu = 1.45 normalised power, CSp = 0.77 normalised power,

Wilcoxon sign rank test p = 0.0002, Delta early cluster 2: CSu = 0.78 normalised power, CSp = 0.31 normalised power, Wilcoxon sign rank test p = 0.0075, Sigma early cluster 1: CSu = 0.36 normalised power, CSp = 0.23 normalised power, Wilcoxon sign rank test p = 0.05).

To provide more information regarding individual difference and the contribution of each participant to the results we calculated individual EEG power in each cluster in all frequency bands. Specifically, we calculate the difference in EEG power between CSu and CSp (Figure S5 ad-e), and the difference in EEG power between early- and late training in the CSu-CSp difference (Figure S5, f-g). We then computed (a) the percentage of participants in the same direction as the effect (Unpleasant > Pleasant or Early > Late training), and (b) the ratio between EEG power in the direction of effect and in the opposite direction (Unpleasant > Pleasant)/(Pleasant > Unpleasant or Early > Late)/(late > Early). We added this information to Figure S5.

For the relation between learning-related neural and behavioural responses analysis, two participants lacked the retention paradigm (morning testing) due to technical error and one participant's sniff response was an outlier (> 3 SD). Excluding these three participants, data from 35 participants remained for the regression analysis in ealry-training and 33 in late-training. This sample size is relatively moderate when examining correlations, and therefore should be taken into account when interpreting the relationship between neural activity during sleep and behaviour in the following morning.

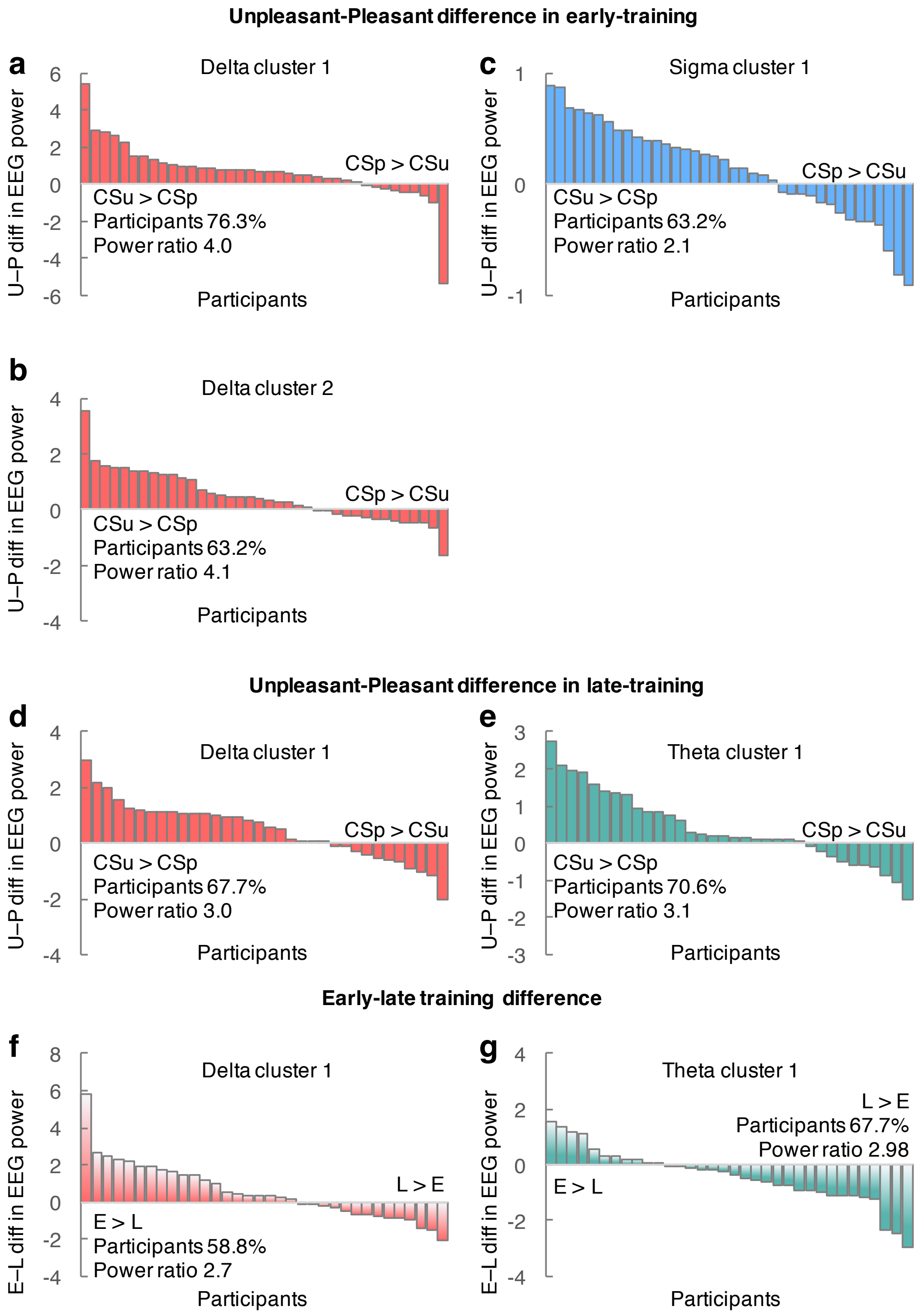


**Figure S5: Individual EEG power**

**(a-c)** EEG power difference between CSu and CSp in early-training in **(a)** delta cluster 1, **(b)** delta cluster 2, and sigma cluster 1. **(d-e)**, EEG power difference between CSu and CSp in late training in **(d)** delta and **(e)** theta. **(f-g)** EEG power CSu-CSP difference between early- and late-training in **(f)** delta and **(f)** theta.

**Respiration related EEG activity**

Respiratory cycle can influence EEG activity where, inhalation and exhalation induce distinct EEG patterns (Chervin et al. 2004; Perl et al. 2019). In our study, we compare EEG power between two conditions with different sniff volume: CSu induced smaller sniff volume than CSp. To test whether differences in sniff volume between conditions could account for the observed changes in EEG power we examined whether inhalation magnitude modulate EEG power. To disentangle the influence of sniff magnitude and learning, and keep the same sleep structure, we analysed the inhalation occurring before tone onset. We time-locked the EEG activity to the onset of inhalation and divided the trials in small (below median) and large (above median) inhalation volume for each participant. First, we verified that inhalation volume indeed differed between the small and large inhalation condition ($t_{37}$ = 10.5 $p < 0.00001$). Next, using cluster permutation test we compare EEG power in delta, theta and sigma between small and large inhalation conditions. We found no significant cluster in either delta, theta or sigma across conditions (all $p_{cluster} > 0.15$). This finding suggests that the observed EEG power differences between CSp and CSu did not result from differences in inhalation volume.
